# Environmental drivers of disease depend on host community context

**DOI:** 10.1101/2021.01.26.428219

**Authors:** Fletcher W. Halliday, Mikko Jalo, Anna-Liisa Laine

## Abstract

Predicting disease risk in an era of unprecedented biodiversity and climate change is more challenging than ever, largely because when and where hosts are at greatest risk of becoming infected depends on complex relationships between hosts, parasites, and the environment. Theory predicts that host species characterized by fast-paced life-history strategies are more susceptible to infection and contribute more to transmission than their slow-paced counterparts. Hence, disease risk should increase as host community structure becomes increasingly dominated by fast-paced hosts. Theory also suggests that environmental gradients can alter disease risk, both directly, due to abiotic constraints on parasite replication and growth, and indirectly, by changing host community structure. What is more poorly understood, however, is whether environmental gradients can also alter the effect of host community structure on disease risk. We addressed these questions using a detailed survey of host communities and infection severity along a 1100m elevational gradient in southeastern Switzerland. Consistent with prior studies, increasing elevation directly reduced infection severity, which we attribute to abiotic constraints, and indirectly reduced infection severity via changes in host richness, which we attribute to encounter reduction. Communities dominated by fast pace-of-life hosts also experienced more disease. Finally, although elevation did not directly influence host community pace-of-life, the relationship between pace-of-life and disease was sensitive to elevation: increasing elevation weakened the relationship between host community pace-of-life and infection severity. This result provides the first field evidence, to our knowledge, that an environmental gradient can alter the effect of host community structure on infection severity.

## Introduction

Human pressure is generating an unprecedented expansion of conditions associated with global change, including the emergence and spread of infectious diseases (Jones *et al.* 2008; Fisher *et al.* 2012; Gibb *et al.* 2020), changing species diversity (Hillebrand *et al.* 2018; Díaz *et al.* 2019), and a changing climate (Pachauri *et al.* 2014). Understanding how these processes interact with one another is one of the greatest research challenges of the 21^st^ century, and will be the key to limiting their impacts on food production systems, wildlife, and humans. In the context of infectious disease, limiting the impacts of global change requires the ability to predict when and where crops, livestock, wildlife, and humans are at the greatest risk of becoming infected. Disease ecology provides a framework for achieving this goal through careful examination of interactions among hosts, parasites, and the environment (McNew 1960; Johnson *et al.* 2015b; Seabloom *et al.* 2015) (Fig. 1a). Yet we have a poor understanding of how this framework operates under natural conditions, in part because several mechanisms can operate simultaneously, making it difficult to tease apart their relative contributions to realized disease risk.

**Figure 1.**
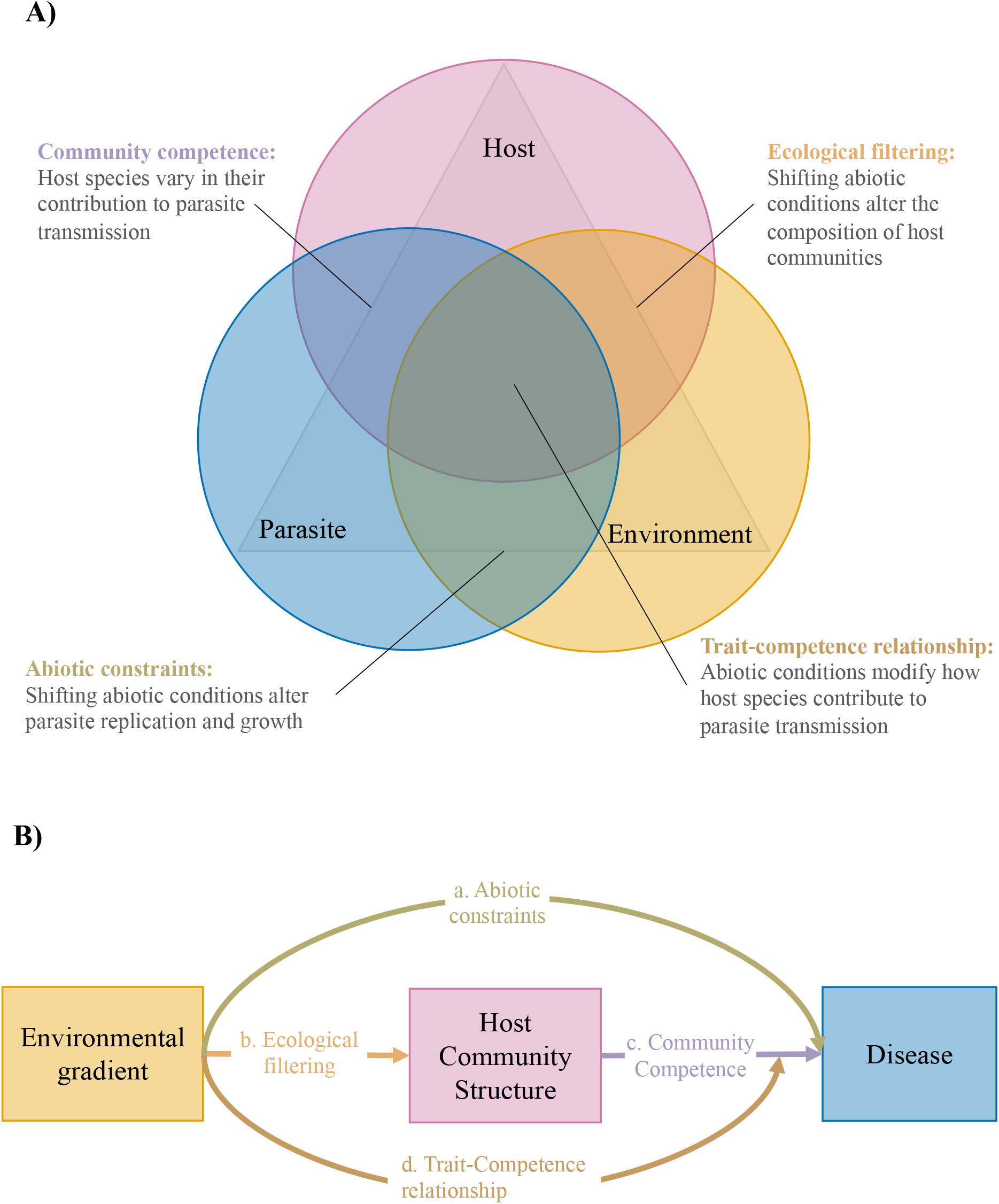
Relationships among hosts parasites and their environment at the scale of host communities. A) The disease triangle (McNew 1960) suggests that a combination of host, parasite, and environmental factors will influence whether disease is observed in a given location. Here, we conceptualize the disease triangle at the community level as consisting of three overlapping or interacting factors to demonstrate how the influence of global change on disease risk might depend on how these factors overlap. B) Conceptual metamodel of an environmental gradient directly influencing disease risk (path a), and indirectly influencing disease risk, both by altering host community structure (paths b and c), and by modifying how host community structure influences disease risk (path d).

Global change drivers such as increased temperature and increased nitrogen deposition can profoundly alter disease risk (Harvell *et al.* 2002; Garrett *et al.* 2006; Rohr *et al.* 2011; Veresoglou *et al.* 2013). These effects can result from direct impacts of global change drivers on parasite growth, survival and reproduction that underpin disease risk. For example, in an experiment in the Rocky mountains, host plants that grew on heated research plots showed increased disease, largely by increasing the amount of time that environmental conditions were favorable for parasite growth and reproduction (Roy *et al.* 2004). Importantly however, these same environmental factors can also indirectly influence disease risk by altering the composition of host or vector communities that are required for sustained parasite transmission (Harvell *et al.* 2002; Garrett *et al.* 2006; Newton *et al.* 2011; Rohr *et al.* 2011; R. Yáñez-López 2012; Elad & Pertot 2014; Mordecai *et al.* 2019). Thus, shifts in parasite replication that are driven by changing host or vector distributions can also determine whether and how global change will alter disease risk. There is growing empirical evidence in support of both direct effects that alter parasite growth and replication, as well as indirect effects that are mediated by changing host or vector community structure. However, disentangling the relative impacts of these direct and indirect effects of global change on disease risk has been historically challenging, because it often requires a-priori knowledge of environmental constraints acting on host and parasite populations (Harvell *et al.* 2002; Garrett *et al.* 2006; Rohr *et al.* 2011; Mordecai *et al.* 2019).

One way to disentangle direct and indirect effects of environmental conditions on disease is to consider these effects in the context of host functional traits. Host functional traits underlie ecologically important resource acquisition and allocation tradeoffs: hosts must balance allocating limited resources to maximize growth and reproduction, while also constructing tissue capable of withstanding stressful environmental conditions (Reich *et al.* 2003a; Wright *et al.* 2004; Reich 2014; Díaz *et al.* 2016). Thus, linking environmental gradients with relevant functional traits has become a tractable way to predict community structure in the face of global change (Diaz & Cabido 1997; Lavorel & Garnier 2002; Cornelissen *et al.* 2003; McMahon *et al.* 2011; Sundqvist *et al.* 2013; Reich 2014; Funk *et al.* 2017; Kattge *et al.* 2020).

The functional traits expressed by those species that are able to colonize and persist in a given location can, in turn, affect disease risk (Johnson *et al.* 2013; Halliday *et al.* 2019; Kirk *et al.* 2019). Specifically, an infected host’s ability to transmit disease to uninfected hosts, a trait often referred to as host competence, is often related to fast-growing, poorly defended tissues and short lifespans (Cronin *et al.* 2010, 2014; Johnson *et al.* 2012; Huang *et al.* 2013; Martin *et al.* 2016, 2019; Becker & Han 2020; Stewart Merrill & Johnson 2020; Welsh *et al.* 2020). Importantly, these functional trait values also underlie ecological tradeoffs related to host growth and defense, resource acquisition and allocation, and life history (Stearns 1989; Herms & Mattson 1992; Stearns 1992; Ricklefs & Wikelski 2002; Reich *et al.* 2003b; Wright *et al.* 2004; Reich 2014; Martin *et al.* 2016). Thus, host community competence is expected to correspond to the same functional traits that link host community structure to global change drivers.

A trait-based framework of host community competence may explain why biodiversity loss is consistently associated with higher disease risk (LoGiudice *et al.* 2003; Ostfeld & LoGiudice 2003; Johnson *et al.* 2013; Halliday *et al.* 2020b), a relationship known as the “dilution effect” of biodiversity (Ostfeld & Keesing 2000; Keesing *et al.* 2006, 2010). This is because host species that are most resistant to biodiversity loss or best able to colonize newly disturbed habitats often rely on the same life-history strategies that are associated with higher host competence (LoGiudice *et al.* 2003; Ostfeld & LoGiudice 2003; Johnson *et al.* 2013). For example, species that are associated with habitat fragmentation, a key anthropogenic driver of biodiversity loss, are often characterized by life history strategies favoring a “fast pace of life” (i.e., fast growth rates, quick reproduction, and high dispersal) (Hanski *et al.* 2006; Gibbs & Van Dyck 2010; Albrecht & Haider 2013; Fay *et al.* 2015; Keinath *et al.* 2017; Merckx *et al.* 2018; Ziv & Davidowitz 2019). But this fast pace of life often comes at the cost of reduced defense against parasites (Herms & Mattson 1992; Cronin *et al.* 2010, 2014; Johnson *et al.* 2012; Sears *et al.* 2015; Heckman *et al.* 2019; Cappelli *et al.* 2020). Thus, habitat fragmentation can increase disease by increasing the density of fast pace-of-life, highly competent hosts, while slow-pace-of-life, less-competent hosts are lost (Joseph *et al.* 2013; Mihaljevic *et al.* 2014; Johnson *et al.* 2015a). This hypothesis has widespread empirical support in a variety of systems (Ostfeld & LoGiudice 2003; Johnson *et al.* 2013, 2019; Liu *et al.* 2018). Shifting community structure during biodiversity loss may therefore predictably influence infectious disease risk (Halliday *et al.* 2020b).

Although relationships between host community structure and disease risk are becoming increasingly appreciated, how these relationships change across environmental gradients and their consequences for disease risk under global change remain unknown (Halliday & Rohr 2019; Halliday *et al.* 2020b). The relationship between host traits and host competence can be variable, and this relationship might also depend on the environmental context in which host-parasite interactions play out (Fig 1b., path d). For example, Welsh et al (2016) showed that when hosts were reared under novel resource conditions, trait-based models of host susceptibility became increasingly inaccurate, because novel resource conditions altered how traits covaried with one another and how raw trait values predicted infection. Thus, traits associated with host community competence in one environment might not predict host community competence under environmental change.

We hypothesized that three non-mutually exclusive mechanisms would determine how global change influences disease risk in host communities: (1) **directly, by altering parasite growth and reproduction**(i.e., through abiotic constraints; Fig 1b., path a), (2) **indirectly, by altering which host species occur in which locations**(i.e, through shifting host community structure; Fig 1b., paths b and c), and (3) **interactively, by altering how host traits influence parasite transmission**(i.e., by altering the relationship between host traits and host competence; Fig 1b., path d).

Here, we test the relative contributions of these three mechanisms through which environmental conditions can drive infectious disease risk by measuring foliar fungal disease in host plant communities along a roughly 1100 m elevational gradient in Southeastern Switzerland. Foliar fungal parasites are a widely used, tractable model of disease risk that respond to small-scale variation in host community structure and environmental conditions (Mitchell *et al.* 2002, 2003; Rottstock *et al.* 2014; Liu *et al.* 2016; Halliday *et al.* 2017; Liu *et al.* 2017, 2018; Halliday *et al.* 2019; Cappelli *et al.* 2020). Host community structure and environmental conditions, in turn, vary predictably with elevation (Grinnell 1914; Whittaker 1956; Malhi *et al.* 2010; Sundqvist *et al.* 2013; Halbritter *et al.* 2018). Thus, an elevational gradient represents a natural laboratory for studying long-term, large scale changes in climate as well as interacting biotic and abiotic factors that are associated with global change (Fukami & Wardle 2005; Sundqvist *et al.* 2013; Alexander *et al.* 2015).

Our study reveals strong evidence that elevation can directly influence disease risk, which we attribute to abiotic constraints on parasite replication and growth, can indirectly influence disease risk by shifting host community structure, and can interactively influence disease risk by modifying the trait-competence relationship. Furthermore, the association between host community structure and disease was as strong as the combined effects of elevation on disease risk. Together, these results highlight the need to consider biotic and abiotic drivers jointly, in order to predict disease risk in the face of global change.

## Methods

### Study system

To evaluate abiotic constraints on parasite replication and growth, shifting host community structure, and modification of the trait-competence relationship as mechanisms through which environmental gradients can influence disease risk, we surveyed 220, 0.5 m-diameter vegetation communities, that were established as part of the Calanda Biodiversity Observatory (CBO) in 2019 in order to investigate biotic and abiotic drivers of species interactions. The CBO consists of four publicly owned meadows located along a 1101 m elevational gradient (648 m to 1749 m) below tree-line on the south-eastern slope of Mount Calanda (46°53′59.5″N 9°28′02.5″E) in the canton of Graubünden (Fig. 2). The mean annual temperature at 550 m altitude is 10 °C and the mean annual precipitation is 849 mm (MeteoSwiss 2020), with temperature declining and precipitation increasing as elevation increases (e.g., in 2013 and 2014, mean temp and precipitation at 1400 m were 7 °C and 1169 mm, respectively; Alexander *et al.* 2015). The soil in the area is generally calcareous and has low water retention (Eggenberg & Möhl 2007; Alexander *et al.* 2015). The four CBO meadows are variable in size (roughly 8 to 40 Ha), and separated by forests and at least 500 m elevation. Meadows are maintained through grazing and mowing, a typical form of land use in the Swiss Alps (Bätzing 2015), and cover collinean (<800 m) mountain (800 m–1500 m) and subalpine (1500–2200 m) vegetation zones (Ozenda 1985; Eggenberg & Möhl 2007). The CBO meadows are grazed by cattle twice per year as the cattle are moved between low and high altitudes.

**Figure 2.**
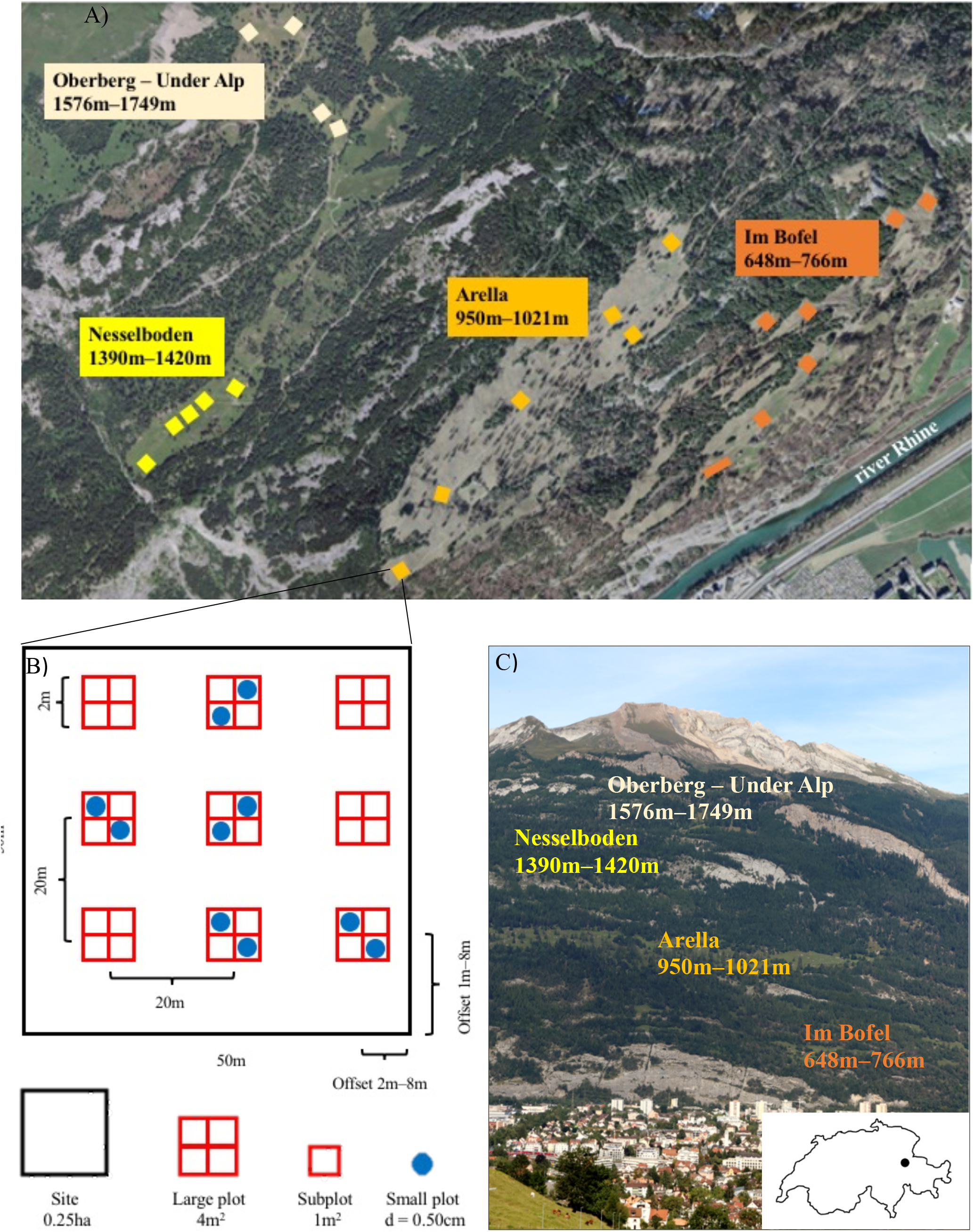
Overview of the Calanda Biodiverstity Project. A) Study meadows and sites on mount Calanda. Photo: Federal Office of Topography SwissTopo 2020, editing: Mikko Jalo B) Example of the arrangement of large and small plots within a site. C) The study meadows on mount Calanda. Photo and editing: Mikko Jalo

The elevational gradient captured by the CBO allows us to explore associations among abiotic factors and biodiversity while minimizing other confounding factors like day length, geology and biogeographic history (Halbritter *et al.* 2018). Increasing elevation is associated with changes in a variety of abiotic conditions, including a reduction in temperature. Temperature decreases approximately 0.4 −0.7 °C for each 100 m increase in elevation because of lower air pressure in high elevations, a phenomenon known as the altitudinal temperature lapse rate (Barry 2008). The altitudinal temperature lapse rate varies among years and even days, usually being lower in winters and during nights. Typical altitudinal temperature lapse rates in the Alps vary from −0.54 °C/100 m to −0.58 °C/100 m (Rolland 2003).

### Study design

The CBO consists of a nested set of observational units (Fig. 2). Each meadow contains 4–7, ․25 ha sites (n = 22 sites). Sites were selected to maximize coverage over each meadow, avoiding roads that would cross the sites and large trees, shrubs and rocks that could create a forest- or shrub-type habitat that differs from grassland. Each site is 50m × 50m and contains a grid of nine evenly spaced, 4m^2^ large-plots, with the exception of one site (I3), which is 100 m × 25 m and contains 10 large plots due to its shape. Altogether, there are 199 large plots. In each site, large plots are arranged in a grid with each plot separated by at least 20m distance from its nearest neighbor. The location of the grid was randomized within each site and always located at least 2m from the site edge. Each large plot is subdivided into four, 1 m^2^ subplots (n = 796). At each site, five large plots were selected to contain an intensively surveyed module (ISM), which consisted of two 50 cm-diameter, round small plots, placed in opposite subplots (n=110 ISMs consisting of 220 small plots). These intensively surveyed small plots are the smallest unit of observation used in this study (Fig. 2).

### Quantification of host community structure

In July 2019, we recorded the identity and visually quantified the percent cover of all plant taxa in each small plot (n = 220). Vegetation surveys entailed the same two researchers searching within the subplot area for all vascular plants present in the subplot, before jointly estimating the total percent cover of each species (Halbritter *et al.* 2020). Plant individuals that were growing outside the small plot, but whose foliage extended into the small plot, were included in this survey. Plant taxa were identified with the help of plant identification literature (Eggenberg & Möhl 2007; Eggenberg *et al.* 2018; Lauber *et al.* 2018). The survey started at the lowest elevation and continued higher in order to survey the meadows approximately at the same phase of the growing season in relation to one another. The survey was initiated at least four days after cows had grazed each meadow (Table S1).

We evaluated changes in three components of host community structure to evaluate indirect effects of elevation on disease: host species richness, community-weighted mean host pace of life, and richness-independent phylogenetic diversity of host species. We quantified community-weighted mean host pace of life using the TRY database (Kattge *et al.* 2020). We first extracted six traits for every host taxon in the database (Plant photosynthetic rate, leaf chlorophyll content, leaf lifespan, leaf nitrogen content, leaf phosphorus content, and specific leaf area), omitting tree seedlings, which are functionally dissimilar from the more dominant herbaceous taxa, and taxa that could not be identified to host genus, which together, never accounted for more than 7% cover in a plot (mean = 0.04%). Unknown taxa that could be identified to the genus level were assigned genus-level estimates for each host trait, by taking the mean of the trait value for all members of that genus that had been observed on Mount Calanda during extensive vegetation surveys (Table S2). We then performed full-information maximum-likelihood factor analysis to produce a single axis representing covariation in the functional traits associated with host pace-of-life using the umxEFA function in r-package umx (Bates *et al.* 2019). This approach allows each host taxon to be assigned a value for host pace-of-life, even if that taxon is missing some values for individual functional traits. Finally, we calculated a single value for each small plot (n = 220) using the community-weighted mean of host pace-of-life (hereafter community pace of life). The community weighted mean (CWM) was calculated as:

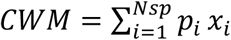

where Nsp is the number of taxa within a plot with a pace-of-life trait value in the dataset, p_i_ is the relative abundance of taxon, *i*, in the plot, and x_i_ is the host pace-of-life value for taxon, *i*.

To quantify phylogenetic diversity, independent of species richness, a phylogeny of all non-tree species was constructed using the phylo.maker function in R package V.PhyloMaker (Jin & Qian 2019). Plant phylogenetic diversity for each small plot was calculated using the ses.mpd function in R package Picante (Kembel *et al.* 2010). This function uses a null-modeling approach that measures the degree to which a small plot is more or less phylogenetically diverse than random, given the number of host species, weighted by their relative abundance. Specifically, we generated a z-score that compares the mean-pairwise-phylogenetic-distance between taxa in a plot to a randomly assembled plot with the same number and relative abundance of host species, drawn from the full species pool, and permuted 1000 times.

### Quantification of disease

A survey of foliar disease symptoms was carried out in August 2019 by estimating the percent of leaf area damaged by foliar fungal parasites on up to five leaves of twenty randomly selected host individuals per small plot (n = 18203 leaves on 4400 host individuals across 220 small plots). The disease survey was conducted by placing a grid of 20 equally spaced grill sticks into the ground, with each stick having a distance of 10 cm to its nearest neighbor (Fig S1). The 20 plant individuals that were most touching the sticks were then identified, and the five oldest non-senescing leaves on each plant were visually surveyed for foliar disease symptoms following the plant pathogen and invertebrate herbivory protocol in Halbritter et al (2020). The survey was carried out on leaves, because symptoms are highly visible and easily grouped into parasite types on leaves. On each leaf, we estimated the leaf area (%) that was covered by disease symptoms. Some plant individuals had fewer than five leaves, so fewer than five leaves were surveyed on those plants. Unlike the vegetation survey, the disease survey was not conducted in elevational order due to logistical constrains related to site accessibility. Small plots were surveyed between 29 July and 19 August 2019 (Table S1), which we observed to be time of peak plant biomass in this system.

Disease was assessed for each small plot using community parasite load, calculated as the mean leaf area damaged by all parasites on a host, multiplied by the relative abundance of that host, and then summed across all hosts in the plot (e.g., Mitchell *et al.* 2002; Halliday *et al.* 2017, 2019).

### Quantification of abiotic conditions

Soil temperature (6 cm below the soil surface), soil surface temperature, air temperature (12 cm above the soil surface), and soil volumetric moisture content were recorded at 15 minute intervals for 22–37 days (average 31 days) in the central large plot of each site (n = 22) using a TOMST-4 datalogger (Wild *et al.* 2019) (Fig. S1). The total duration of measurement varied because some of the dataloggers had to be moved earlier or temporarily because of mowing or grazing activities (Table S1).

### Statistical analysis

All statistical analyses were performed in R version 3.5.2 (R Core Team 2015) and consisted of fitting linear mixed models with an identity link and Gaussian likelihoods using the lme function in the nlme package (Pinheiro *et al.* 2016). In order to meet assumptions of normality and homoscedasticity, we square-root transformed community parasite load and added an identity variance structure (varIdent function) for each site, which based on visual inspection of residuals of each model, exhibited considerable heteroscedasticity (Zuur *et al.* 2009; Pinheiro *et al.* 2016). Each model included large plots, sites, and meadows as nested random intercepts to account for non-independence among observations due to the sampling design of the CBO.

We first explored the relationship between each measure of host community structure (i.e., host species richness, richness-independent phylogenetic diversity, and host community pace of life) and elevation by constructing three models each consisting of one measure of host community structure as a response variable and elevation as the only fixed effect.

We next tested whether the relationship between host community structure and disease would change as a function of elevation by constructing a mixed model with square-root transformed community parasite load as the response, and elevation, host community species richness, pace of life, and richness-independent phylogenetic diversity as fixed effects. To estimate whether the effect of host community structure depends on elevation, we also included in the model the pairwise interactions between each of the three measures of host community structure and elevation as additional fixed effects. To evaluate model fit, we calculated the root-mean-squared error (RMSE) of the model, the marginal and conditional pseudo-R^2^ of the model using the r.squaredGLMM function in the MuMIn package (Bartoń 2018), and the RMSE using leave-one-out cross validation (LOOCV RMSE). To test whether effects driven by host community pace of life were influenced by one or a few important functional traits, we repeated this analysis, including the standardized and centered community-weighted-mean of each leaf trait (leaf chlorophyll content, leaf lifespan, leaf nitrogen content, leaf phosphorus content, and specific leaf area) replacing host community pace of life. However, none of these models were improvements over the model including host community pace of life (Table S3), and were therefore excluded from further analyses.

Finally, to compare direct effects of elevation on disease risk with indirect effects of elevation that are mediated by changing host community structure, we performed confirmatory path analysis using the PiecewiseSEM package (Lefcheck 2016). Specifically, we fit a structural equation model (SEM) that included a direct effect of elevation on square-root-transformed disease, the effect of elevation on three endogenous mediators (host community species richness, pace of life, and phylogenetic diversity, which together measure changes in host community structure (following Halliday *et al.* 2019, 2020a), and the effects of those three mediators on square-root-transformed community parasite load. We also tested the hypothesis that elevation altered the relationship between host community structure and disease by fitting a second-stage moderated mediation (Hayes 2015) including the pairwise interaction between elevation and community pace of life, omitting other potential interactions that were non-significant in the model testing whether effects of community structure on disease depend on elevation.

## Results

### Association between elevation and abiotic factors

Mean soil, soil surface, and air temperature strongly and consistently decreased with increasing elevation (r = −0.94; r = −0.93; r = −0.95; respectively), while mean soil moisture was uncorrelated with elevation (r = −0.07). The mean soil surface temperature at sites located in the highest elevation meadow (1576 m–1749 m) was, on average, 4.67 °C (26 %) lower than sites located in the lowest elevation meadow (648 m–766 m). The altitudinal temperature lapse rate along the elevational gradient was −0.57 °C/100 m. Because temperature was collinear with elevation and soil moisture was uncorrelated with elevation, both abiotic factors were omitted from further analyses.

### Effect of elevation on host community structure

In total, 188 host taxa were observed across the 220 small plots of the CBO. The communities consisted mostly of perennial herbs such as *Salvia pratensis* and *Helianthemum nummularium*, and were dominated by grasses that tolerate grazing such as *Dactylis glomerata*, *Lolium perenne* and *Phleum pratense*. The most abundant species was *Brachypodium pinnatum*. An herbarium specimen of each taxon encountered is deposited at the University of Zürich. Species richness in the small plots varied from 7–30 species (median 20), and increased with increasing elevation (p = 0.044; Marginal R^2^ = 0.07; Conditional R^2^ = 0.75), with median species richness roughly 16 % higher in plots located at the highest elevation meadow compared to the lowest elevation meadow (Table S4).

We performed confirmatory factor analysis to assign six foliar functional traits associated with the worldwide leaf economics spectrum to a single axis representing host pace-of-life. One trait, photosynthetic rate, loaded particularly poorly on this axis (factor loading 0.036), and was therefore excluded from the latent factor. This resulted in a single factor, explaining 62% of the variance in specific leaf area, 51% of the variance in leaf chlorophyll content, 25% of the variance in leaf nitrogen, 10% of the variance in leaf phosphorus, and 2% of the variance in leaf lifespan (χ² (df=5) = 4.24, p = 0.52; CFI = 1.019; Figure S2). Consistent with the resource-acquisition and allocation tradeoffs, (Wright *et al.* 2004; Reich 2014; Díaz *et al.* 2016), higher values of host pace of life were associated with increases in specific leaf area, leaf chlorophyll content, leaf nitrogen, and leaf phosphorus, and with shorter leaf lifespans. In contrast with the expectation that increasing elevation would reduce host community pace of life via shifting host community structure (Hulshof *et al.* 2013; Descombes *et al.* 2017; but see Pellissier *et al.* 2018), host community pace of life was only marginally influenced by elevation (p = 0.069; Marginal R^2^ = 0.11; Conditional R^2^ = 0.83; Table S4; Fig S3).

Host phylogenetic diversity was calculated independent of host species richness, by computing the degree to which host communities were more or less phylogenetically diverse than random, given the number of host species and their relative abundance. Across all sites, host communities were commonly phylogenetically clustered, meaning that host species were more closely related than would be expected based on the number of host species and their relative abundance (Fig S3). Host phylogenetic diversity was independent of host species richness (r = 0.17) and increased (i.e., became less phylogenetically clustered) with increasing elevation (p = 0.019; Marginal R^2^ = 0.12; Conditional R^2^ = 0.71; Table S4; Fig S3).

### Model testing effects of elevation, community structure, and their interaction on disease

Our mixed model of disease revealed several independent and interactive effects of host community structure and elevation on disease risk (Marginal R^2^ = 0.238; Conditional R^2^ = 0.460; RMSE = 0.295; LOOCV RMSE = 0.315; Table S5). Consistent with the hypothesis that host pace-of-life can determine host community competence, communities that were dominated by hosts with fast-paced life-history strategies exhibited the most disease, but this effect declined as elevation increased (elevation × pace-of-life: p = 0.0016). This weakening effect of host community pace of life with increasing elevation is consistent with the hypothesis that elevation can alter which traits favor parasite transmission through the relationship between host competence and disease risk (Fig. 3). These results therefore provide field evidence that an environmental gradient can alter the effect of host community structure on disease risk.

**Figure 3.**
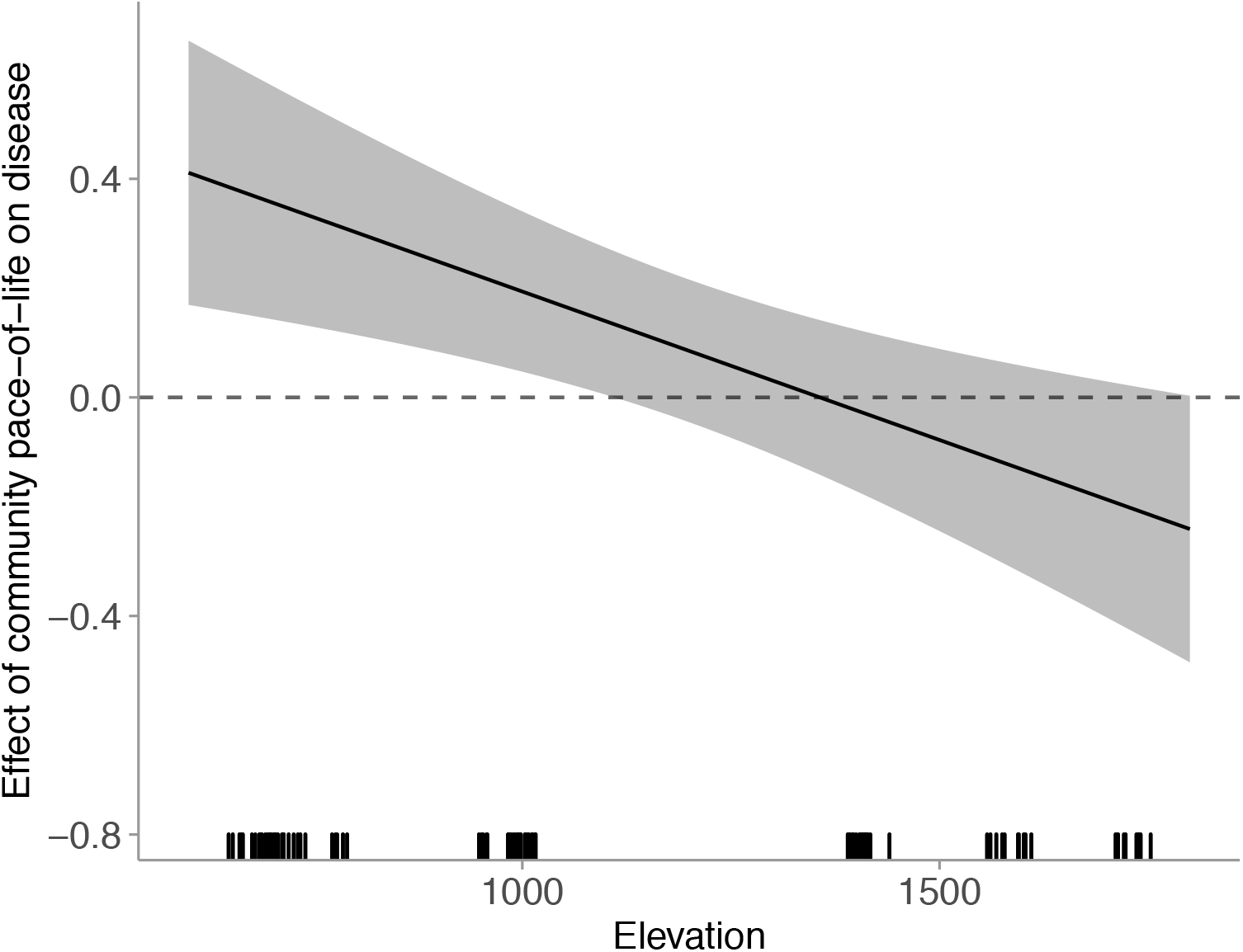
Effect of host community pace of life on disease as a function of increasing elevation. Model estimated effects of elevation on the slope of the relationship between host community pace of life and (square-root-transformed) parasite community load (i.e., the interactive effect of host community pace of life and elevation on disease). The rug along the *x*‐axis shows the distribution of the empirical data. Communities that are located at the lowest elevation exhibit the strongest positive relationship between host pace of life and disease. That positive relationship weakens as elevation increases, and above 1000m, there is no relationship between host pace of life and disease.

The model also revealed significant independent effects of host community structure and elevation on disease risk. Specifically, the model revealed evidence supporting the dilution effect hypothesis: increasing species richness was associated with a reduction in disease (p = 0.0057), and this effect was independent of elevation (elevation × richness: p = 0.21). Community parasite load was also negatively associated with increasing elevation (p = 0.0077), consistent with the hypothesis that elevation can alter parasite growth and reproduction via abiotic constraints. In contrast, there was no relationship between host phylogenetic diversity and disease (p = 0.66), nor was there an interaction between phylogenetic diversity and elevation on disease (p = 0.32).

### Structural equation model comparing direct and indirect effects of elevation on disease

Together, models of host community species richness, pace of life, and phylogenetic diversity showed that elevation could determine changes in host community structure, and the model of disease showed that host community structure and elevation could independently and interactively influence disease. To explore the relative influence of these direct, indirect, and interactive effects on disease risk, we next constructed a structural equation model. Our data were well fit by this model (Fisher’s C = 1.463; P-value = 0.481; 2 degrees of freedom, Table S6). The model compares three separate pathways through which increasing elevation reduced disease: First, elevation reduced community parasite load directly (standardized path coefficient = −0.23). Second, elevation reduced community parasite load indirectly by increasing host species richness (product of standardized path coefficients = 0.038). Third, increasing elevation reduced community parasite load interactively by weakening the relationship between host pace of life and disease risk (Standardized path coefficient = −0.61; Fig. 4a). Yet, despite the strong and consistent reduction in disease risk with increasing elevation, this effect was still roughly equivalent to the direct effects of host community structure on disease (Sum of absolute value of standardized path coefficients = 0.82; Fig. 4b). Furthermore, host community pace of life was still the single strongest predictor of disease risk in this study (standardized path coefficient = 0.66). Together these results highlight the pressing need to consider host community context in predicting how shifting environmental gradients will alter disease risk.

**Figure 4.**
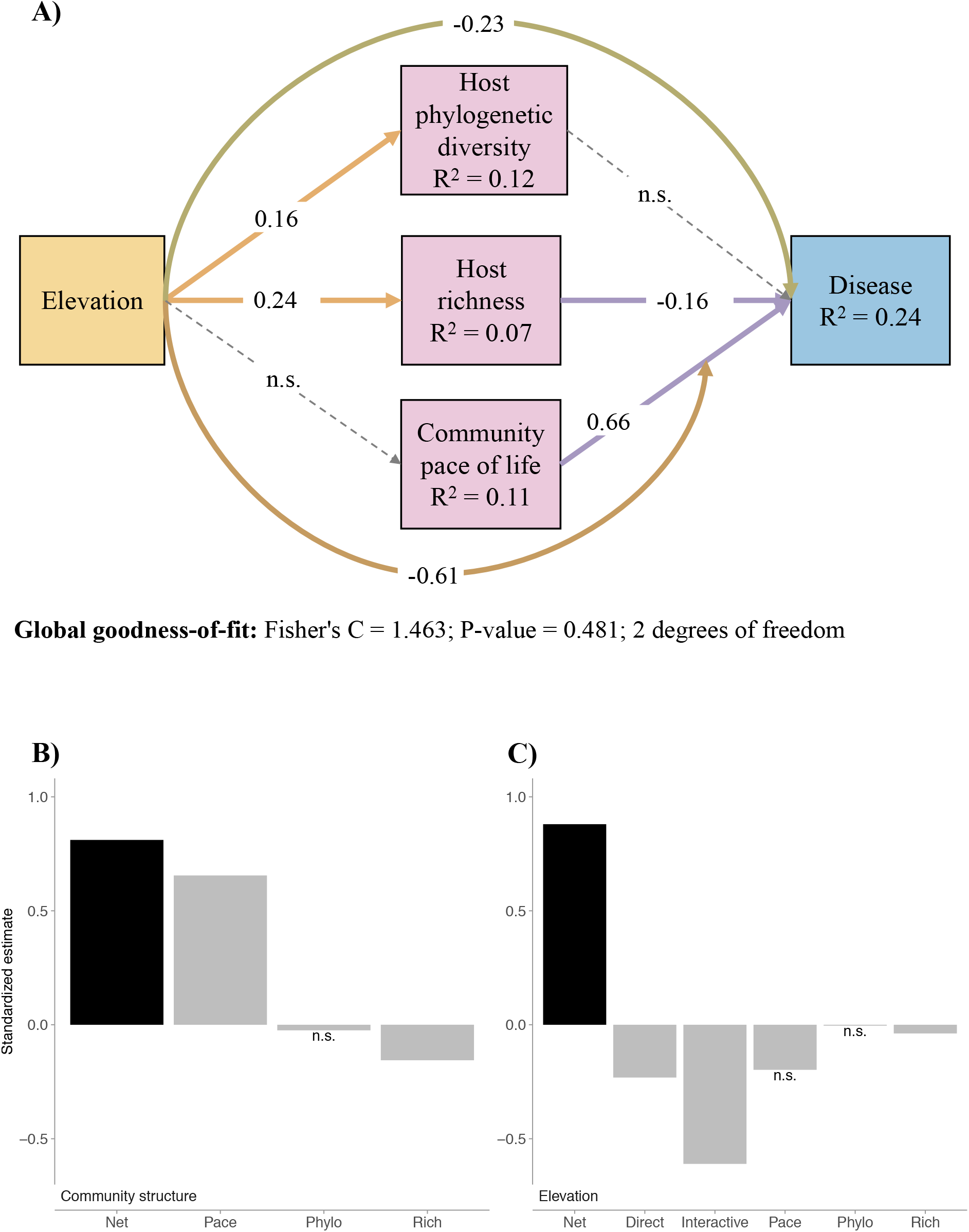
Results from the piecewise structural equation model. A) The structural equation model. Dashed lines are not supported by the model (*p* > ․05). All coefficients are standardized. Correlations between errors were not supported by the model and are not shown. Colors are drawn to match the diagram in Fig. 1 B) Summary of standardized path coefficients associated with host community structure. Grey bars are the effects of three measures of host community structure on parasite community load, representing host community competence as illustrated in Fig. 1 path c, (Pace = host community pace of life; Phylo = host phylogenetic diversity; Rich = host species richness). C) Summary of standardized path coefficients associated with elevation. “Direct” is the direct effect of elevation on community parasite load, representing abiotic constraints as illustrated in Fig. 1 path a. “Interactive” is the statistical interaction between elevation and host pace of life, representing the trait-competence relationship illustrated in Fig. 1 path d. The pace, phylo, and rich bars are the product of indirect pathways involving host pace of life, host phylogenetic diversity, and host species richness, respectively, together representing the indirect effect driven by shifting host community structure and host community competence illustrated in Fig. 1 paths b and c. In both B) and C), paths that were not supported in the model are annotated with “n.s.”. The net effect (shown in black) is quantified as the sum of the absolute value of all paths, excluding those that were not supported in the model. Higher elevation reduced disease through three non-mutually exclusive pathways: directly via abiotic constraints, and indirectly both via shifting host community structure as well as by altering the trait-competence relationship.

## Discussion

This study shows, to our knowledge, the first field evidence that, in addition to directly influencing disease risk, the abiotic environment can also indirectly influence disease by altering host community structure and can interactively influence disease by modifying how host community structure influences disease risk. Furthermore, the abiotic environment explained roughly the same amount of variation in disease risk as host community structure, suggesting that any single factor was inadequate for explaining disease risk along our environmental gradient. Together, these results reveal the role that host communities play in protecting ecosystem health across environmental gradients, suggesting that predicting how global change will influence disease risk will require explicit consideration of how host and parasite communities respond to global change.

Our results indicate that increasing elevation can directly influence disease. Specifically, increasing elevation reduced disease, even after accounting for effects of host community structure on disease. We hypothesize that increasing elevation may have reduced disease directly by constraining parasite growth, survival, and reproduction. Many foliar parasites grow and reproduce more successfully in warmer environmental temperatures, which occur more commonly at low elevation (Tapsoba & Wilson 1997; Harvell *et al.* 2002; Waugh *et al.* 2003; Garrett *et al.* 2006; Avenot *et al.* 2017). Warmer temperatures can also increase parasite overwintering success (Burdon & Elmqvist 1996; Pfender & Vollmer 1999) or allow parasites to produce more generations during a longer growing season (Garrett *et al.* 2006). These results corroborate past studies suggesting that environmental gradients can directly alter the strength of biotic interactions (Schemske *et al.* 2009; Pellissier *et al.* 2014; Descombes *et al.* 2017; Roslin *et al.* 2017; Hargreaves *et al.* 2019), including host-parasite interactions (Nunn *et al.* 2005; Abbate & Antonovics 2014; LaManna *et al.* 2017; Allen *et al.* 2020).

In addition to directly influencing disease, our results indicate that increasing elevation can also indirectly influence disease by altering host community structure. Specifically, increasing elevation increased host species richness, which, in turn, reduced disease. The increase in host species richness with increasing elevation might be attributable to the occurrence of both low-elevation and high-elevation adapted species occupying the highest study sites (Colwell & Lees 2000). Communities in the highest elevation meadow, located just below the tree line, included plant species characteristic of low elevations (e.g., *Lathyrus pratensis*, *Lolium perenne* and *Salvia pratensis*) and plant species that tend to occupy high elevation grasslands (*Soldanella alpina*, *Ranunculus montanus* and *Carex sempervirens*), indicating that these high-elevation sites represent an intermediate zone between subalpine and alpine vegetation communities.

Host communities with higher species richness, in turn, experienced less disease (i.e., a dilution effect; Keesing *et al.* 2006, 2010), even after accounting for the direct effects of elevation on disease and other measures of host community structure. Past studies indicate that increasing biodiversity is often associated with a decline in disease risk because host community structure shifts during biodiversity loss to favor more competent hosts (LoGiudice *et al.* 2003; Ostfeld & LoGiudice 2003; Joseph *et al.* 2013; Liu *et al.* 2018; Johnson *et al.* 2019; Rohr *et al.* 2020). However, in contrast with past studies focused on biodiversity loss, our study measured biodiversity change across a natural biodiversity gradient, which is not expected to consistently influence disease risk (Halliday *et al.* 2020b). We hypothesize that increasing species richness may have reduced disease risk in this system by reducing host density (i.e., via encounter reduction; Mitchell *et al.* 2002; Keesing *et al.* 2006). Encounter reduction might be particularly relevant in this system, because, in addition to altering host richness, increasing elevation also influences the length and timing of the growing season, which can affect peak prevalence and the duration of the epidemic season.

Although increasing species richness reduced disease in host communities, this effect was dwarfed by the overwhelming influence of host community pace of life on disease. Because more competent hosts often exhibit fast-paced life history strategies (Cronin *et al.* 2010; Johnson *et al.* 2012; Martin *et al.* 2016; Welsh *et al.* 2020), we expected that host communities dominated by species with a fast pace of life would experience greater disease. Our analysis was consistent with this hypothesis: host community pace of life was the strongest predictor of disease risk across the 1100 m elevational gradient and more than 4x stronger than the effect of host species richness on disease. This result corroborates prior studies, suggesting that the effect of host community structure rather than species richness, *per se*, should most strongly influence disease risk in host communities (Johnson *et al.* 2015a; Parker *et al.* 2015; Young *et al.* 2017; Halliday *et al.* 2019).

In addition to direct and indirect effects, our results further indicate that increasing elevation can interactively influence disease by modifying the effect of host community structure on disease. Specifically, a prior study suggested that the relationship between host traits and host competence might be sensitive to environmental conditions (Welsh *et al.* 2016), which we hypothesized would cause the relationship between host community pace of life and disease risk to shift across environmental gradients. Our analysis was consistent with this hypothesis as well: Even though elevation did not modify host community pace of life, the effect of host community pace of life on disease was sensitive to elevation. Host community pace of life most strongly predicted disease risk at the lowest elevation, but this effect weakened and ultimately disappeared as elevation increased.

These results indicate that increasing elevation can modify the effect of host community pace of life on disease risk, which we attribute to a change in the relationship between host traits and host competence across environmental conditions. However, we cannot rule out the possibility that the interaction between host pace of life and elevation could have been driven by another mechanism, as the values of functional traits expressed by a single species may have changed along the elevational gradient via a phenomenon known as intraspecific trait variation (Messier *et al.* 2010; Albert *et al.* 2011; Violle *et al.* 2012; Funk *et al.* 2017). Studies of functional traits (including this study) typically characterize each species with a single value for each trait, such as the species-level mean, under the assumption that ecologically important traits vary more among species than within species (McGill *et al.* 2006). However, functional traits of individuals within a species can vary due to local adaptation and phenotypic plasticity driven by local context (Messier *et al.* 2010; Albert *et al.* 2011; Violle *et al.* 2012; Funk *et al.* 2017). Thus, intraspecific shifts in the expression of key functional traits during global change could drive the apparent interaction between host community pace of life and elevation. Future studies should explore these mechanisms by directly measuring host and parasite functional traits across environmental gradients like elevation.

Together, the results of this study highlight the need to consider host community context in predicting how global change will alter disease risk. Specifically, in this study, the strongest predictor of disease was host community pace of life, rather than the abiotic environment, and increasing elevation most strongly influenced disease by reducing the magnitude of this effect. These results are consistent with a growing body of literature suggesting that the role of host communities in regulating ecosystem processes is at least partially explained by characteristics of species present in those ecosystems (Mouillot *et al.* 2011; Allan *et al.* 2015; Leitão *et al.* 2016; Van de Peer *et al.* 2018; Bagousse-Pinguet *et al.* 2019; Start & Gilbert 2019; Heilpern *et al.* 2020), but that abiotic factors such as temperature can override the effects of biotic factors on ecosystem processes (Cannone *et al.* 2007; Laiolo *et al.* 2018). These results therefore suggest that predicting how global change will influence disease may depend on complex relationships between global change drivers and the structure of host communities.

## Acknowledgements

We are grateful for insightful suggestions and field assistance from K. Raveala, M. Tiusanen, A. Norberg, I. Kohonen, V. Loaiza, and members of the Laine Lab. This work was supported by the University of Zürich and by grants from the Academy of Finland (296686) to A-LL and the European Research Council (Consolidator Grant RESISTANCE 724508) to A-LL.

## Supplementary materials

**Table S1.** Timing of grazing, vegetation and disease surveys and temperature measurements at each site. Recovery days represents the amount of time between the end of grazing activities and the beginning of the vegetation survey at each site.

| Site | Meadow | Elevation | Date of the vegetation survey | Recovery days | Date of the disease survey | Date of the temperature measurements |
| --- | --- | --- | --- | --- | --- | --- |
| I1 | Im Bofel | 766.3m | 10.7.2019 | 24 | 15.8.2019 | 7.8.–9.9.2019 |
| I2 | Im Bofel | 737.4m | 4.7.2019 | 18 | 13.–<br>15.8.2019 | 7.8.–9.9.2019 |
| I3 | Im Bofel | 711.5m | 3.–4.7.2019 | 17–18 | 29.7.2019 | 7.–28.8.2019 |
| I4 | Im Bofel | 702.4m | 2.–3.7.2019 | 16–17 | 29.–<br>30.7.2019 | 7.–28.8.2019 |
| I5 | Im Bofel | 684.5m | 27.6.2019–<br>2.7.2019 | 11–16 | 18.–<br>30.7.2019 | 7.–28.8.2019 |
| I6 | Im Bofel | 702.2m | 28.6.2019 | 12 | 13.8.2019 | 7.8.–11.9.2019 |
| I7 | Im Bofel | 648.5m | 5.–9.7.2019 | 19–23 | 30.7.2019 | 7.8.–11.9.2019 |
| A1 | Arella | 1020.6m | 16.–<br>17.7.2019 | 17–18 | 8.8.2019 | 7.8.–12.9.2019 |
| A2 | Arella | 984.9m | 16.7.2019 | 17 | 15.–<br>19.8.2019 | 7.8.–12.9.2019 |
| A3 | Arella | 949.8m | 15.–<br>16.7.2019 | 16–17 | 8.8.2019 | 7.8.–11.9.2019 |
| A4 | Arella | 1002.8m | 12.7.2019 | 13 | 22.8.2019 | 7.8.–11.9.2019 |
| A5 | Arella | 1001m | 11.7.2019 | 12 | 7.8.2019 | 7.8.–11.9.2019 |
| A6 | Arella | 981.9m | 10.–<br>11.7.2019 | 11–12 | 7.8.2019 | 7.8.–11.9.2019 |
| N1 | Nesselboden | 1390.3m | 23.7.2019 | 20 | 14.8.2019 | 7.8.–12.9.2019 |
| N2 | Nesselboden | 1405.5m | 22.–<br>23.7.2019 | 19–20 | 2.8.2019 | 7.8.–12.9.2019* |
| N3 | Nesselboden | 1407.7m | 19.–<br>22.7.2019 | 16–19 | 2.–6.8.2019 | 7.8.–12.9.2019* |
| N4 | Nesselboden | 1420m | 19.7.2019 | 16 | 6.–<br>14.8.2019 | 7.8.–12.9.2019* |
| N5 | Nesselboden | 1398.6m | 17.–<br>19.7.2019 | 14–16 | 19.8.2019 | 7.8.–12.9.2019* |
| O3 | Oberberg – Under Alp | 1612.7m | 24.7.2019 | 5 | 31.7.2019 | 7.8.–8.9.2019 |
| O4 | Oberberg – Under Alp | 1576.2m | 23.–<br>24.7.2019 | 4–5 | 31.7.2019 | 7.8.–9.9.2019 |
| U1 | Oberberg – Under Alp | 1745.8m | 25.7.–<br>1.8.2019 | 6–13 | 16.8.2019 | 7.8.–8.9.2019 |
| U2 | Oberberg – Under Alp | 1749.2m | 26.7.–<br>1.8.2019 | 7–13 | 16.8.2019 | 7.8.–8.9.2019 |

\*Data loggers were removed for 12 days 30.8.–10.9.2019 due to escape of cattle from higher elevation meadows.

**Table S2.** Calanda Biodiversity Observatory Vegetation list. This list includes species that were observed during the vegetation survey as well as taxa observed outside of the plots during extensive preliminary surveys of Mount Calanda.

| **Species** | **Habitat preference (Flora Helvetica)** |
| --- | --- |
| *Achillea millefolium* | collin-subalpin (-alpin) |
| *Acinos alpinus* | (montane) subalpine (-alpine) |
| *Acinos arvensis* | kollin-montan (-subalpin) |
| *Aconitum lycoctonum* | kollin-subalpin |
| *Aconitum napellus* | subalpine-alpine |
| *Aegopodium podagraria* | kollin-montan (-subalpin) |
| *Agrimonia eupatoria* | kollin-montan (-subalpin) |
| *Agrostis alpina* | subalpin-alpin |
| *Agrostis capillaris* | kollin-subalpin (-alpin) |
| *Agrostis gigantea* | kollin-subalpin |
| *Agrostis stolonifera* | kollin-subalpin (-alpin) |
| *Agrostis schraderiana* | (montane) subalpine-alpine |
| *Ajuga pyramidalis* | (montane) subalpine-alpine |
| *Ajuga reptans* | kollin-subalpin (-alpin) |
| *Alchemilla conjuncta* | montane-subalpin (-alpin) |
| *Alchemilla xanthochlora* | kollin-alpin |
| *Alchemilla pratensis* | kollin-alpin |
| *Alchemilla nitida* | subalpine (-alpin) |
| *Allium carinatum* | kollin-subalpin |
| *Anacamptis pyramidalis* | Kollin-montan |
| *Androsace chamaejasme* | (montane) subalpine-alpine |
| *Androsace obtusifolia* | (subalpine) alpine |
| *Anemone narcissiflora* | (montane) subalpine (-alpine) |
| *Anthericum ramosum* | kollin-subalpin (-alpin) |
| *Anthoxanthum alpinum* | (montane) subalpine-alpine |
| *Anthoxanthum odoratum* | kollin-alpin |
| *Anthriscus sylvestris* | kollin-subalpin |
| *Anthyllis vulneraria* | kollin-montan (-subalpin) |
| *Aquilegia atrata* | (kollin-) montan-subalpin |
| *Arabis ciliata* | (collin-) subalpine (-alpin) |
| *Arenaria ciliata* | (subalpine) alpine |
| *Arenaria serpyllifolia* | Kollin-montan (-subalpin) |
| *Arnica montana* | (montane) subalpine-alpine |
| *Arrhenatherum elatius* | kollin-montan (-subalpin) |
| *Asperula cynanchica* | kollin-montan (-subalpin) |
| *Aster alpinus* | subalpin-alpin |
| *Aster amellus* | kollin-montan |
| *Aster bellidiastrum* | (kollin-) montan-alpin |
| *Astragalus glycyphyllos* | kollin-montan (-subalpin) |
| *Avenula pubescens* | kollin-subalpin (-alpin) |
| *Bellis perennis* | collin-subalpin (-alpin) |
| *Berberis vulgaris* | kollin-subalpin |
| *Bothriochloa ischaemum* | kollin-montan |
| *Botrychium lunaria* | (kollin-) montan-alpin |
| *Brachypodium pinnatum* | kollin-subalpin |
| *Briza media* | kollin-subalpin (-alpin) |
| *Bromus erectus* | kollin-montan (-subalpin) |
| *Bromus hordeaceus* | kollin-montan (-subalpin) |
| *Bromus inermis* | kollin-montan (-subalpin) |
| *Buphthalmum salicifolium* | kollin-subalpin |
| *Calystegia sepium* | kollin-montan |
| *Campanula cochleariifolia* | (kollin-) montan-alpin |
| *Campanula glomerata* | Kollin-montan (-subalpin) |
| *Campanula patula* | kollin-montan (-subalpin) |
| *Campanula rapunculoides* | kollin-montan (-subalpin) |
| *Campanula rotundifolia* | kollin-subalpin (-alpin) |
| *Campanula scheuchzeri* | (montan-) subalpine-alpine |
| *Carduus defloratus* | montan-alpin |
| *Carduus nutans* | kollin-subalpin |
| *Carex capillaris* | (montane) subalpine-alpine |
| *Carex caryophylla* | kollin-subalpin (-alpin) |
| *Carex flacca* | kollin-subalpin (-alpin) |
| *Carex montana* | kollin-subalpin |
| *Carex ornithopoda* | kollin-subalpin (-alpin) |
| *Carex parviflora* | subalpine-alpine |
| *Carex sempervirens* | (montane) subalpine-alpine |
| *Carlina acaulis* | (collinous) montane-subalpin |
| *Carlina vulgaris* | Kollin-montan |
| *Carum carvi* | kollin-subalpin (-alpin) |
| *Centaurea jacea* | kollin-subalpin |
| *Centaurea montana* | montane-subalpine |
| *Centaurea scabiosa* | kollin-montan (-subalpin) |
| *Centaureum erythraea* | kollin-montan |
| *Cephalanthera longifolia* | kollin-montan |
| *Cerastium alpinum* | (subalpine) alpine |
| *Cerastium fontanum* | kollin-montan (-subalpin) |
| *Chenopodium album* | kollin-subalpin |
| *Chenopodium bonus-henricus* | (collin-) montan-subalpin (-alpin) |
| *Cichorium intybus* | kollin-montan (-subalpin) |
| *Cirsium acaule* | (collin) montane-subalpin (-alpin) |
| *Cirsium arvense* | kollin-montan (-subalpin) |
| *Cirsium spinosissimum* | subalpine-alpine |
| *Clinopodium vulgare* | kollin-subalpin (-alpin) |
| *Colchicum autumnale* | kollin-subalpin |
| *Conyza canadensis* | kollin-montan (-subalpin) |
| *Crepis biennis* | Kollin-montan (-subalpin) |
| *Crocus albiflorus* | montan-alpin |
| *Cynosurus cristatus* | kollin-subalpin |
| *Dactylis glomerata* | kollin-subalpin (-alpin) |
| *Danthonia decumbens* | (kollin-) montane-subalpine |
| *Daucus carota* | kollin-montan (-subalpin) |
| *Deschampsia cespitosa* | kollin-alpin |
| *Dianthus superbus* | (collin) subalpin (-alpin) |
| *Dianthus sylvestris* | kollin-subalpin (-alpin) |
| *Digitaria sanguinalis* | kollin-montan |
| *Dryas octopetala* | subalpin-alpin |
| *Echinochloa crus-galli* | kollin-montan |
| *Echium vulgare* | kollin-subalpin |
| *Elyna myosuroides* | subalpine-alpine |
| *Erica carnea* | (kollin-) montane-subalpine (-alpin) |
| *Erigeron neglectus* | (subalpine) alpine |
| *Erodium cicutarium* | kollin-montan (-subalpin) |
| *Euphorbia cyparissias* | kollin-alpin |
| *Euphrasia minima* | subalpine-alpine |
| *Euphrasia rostkoviana montana* | kollin-alpin |
| *Euphrasia salisburgensis* | kollin-alpin |
| *Festuca arundinacea* | kollin-montan (-subalpin) |
| *Festuca ovina* | kollin-alpin |
| *Festuca quadriflora* | subalpine-alpine |
| *Festuca rubra* | kollin-alpin |
| *Filipendula ulmaria* | kollin-subalpin |
| *Fragaria vesca* | kollin-subalpin |
| *Gagea fragifera* | subalpin-alpin |
| *Galium mollugo* | kollin-montan |
| *Galium verum* | kollin-montan (-subalpin) |
| *Gentiana acaulis* | (montane) subalpine-alpine |
| *Gentiana brachyphylla* | alpine |
| *Gentiana campestris* | montan-subalpin (-alpin) |
| *Gentiana clusii* | (montane) subalpine-alpine |
| *Gentiana verna* | montane-alpine |
| *Geranium columbinum* | kollin-montan |
| *Geranium pyrenaicum* | kollin-montan (-subalpin) |
| *Geranium robertianum* | kollin-montan (-subalpin) |
| *Geranium sylvaticum* | (collinous) montane-subalpin (-alpin) |
| *Geum montanum* | (montane) subalpine-alpine |
| *Globularia nudicaulis* | (montane) subalpine (-alpine) |
| *Gymnadenia conopsea* | kollin-subalpin (-alpin) |
| *Helianthemum alpestre* | (montane) subalpine-alpine |
| *Helianthemum nummularium* | kollin-alpin |
| *Helictotrichon pubescens* | kollin-subalpin (-alpin) |
| *Helictotrichon versicolor* | (montane) subalpine-alpine |
| *Helictotrichon pubescens* | kollin-subalpin (-alpin) |
| *Hepatica nobilis* | kollin-montan (-subalpin) |
| *Hieracium lactucella* | kollin-subalpin |
| *Hieracium pilosella* | kollin-alpin |
| *Hippocrepis comosa* | kollin-subalpin (-alpin) |
| *Homogyne alpina* | (montane) subalpine-alpine |
| *Hordeum murinum* | kollin-montan |
| *Hypericum maculatum* | Montan-alpin |
| *Hypericum perforatum* | kollin-montan (-subalpin) |
| *Knautia arvensis* | kollin-montan (-subalpin) |
| *Koeleria pyramidata* | kollin-subalpin |
| *Lamium maculatum* | kollin-subalpin (-alpin) |
| *Larix decidua* | subalpine |
| *Laserpitium siler* | (kollin-) montan-subalpin |
| *Lathyrus niger* | kollin (-montan) |
| *Lathyrus pratensis* | kollin-montan |
| *Lathyrus vernus* | kollin-montan |
| *Leontodon hispidus* | kollin-alpin |
| *Leucanthemum vulgare* | kollin-subalpin (-alpin) |
| *Ligustrum vulgare* | kollin (-montan) |
| *Linum catharticum* | kollin-subalpin |
| *Lithospermum arvense* | kollin-montan (-subalpin) |
| *Lolium perenne* | kollin-montan (-subalpin) |
| *Lotus corniculatus* | kollin-subalpin (-alpin) |
| *Lotus maritimus* | collin-montan (-subalpin) |
| *Luzula campestris* | kollin-montan (-subalpin) |
| *Luzula nivea* | (collinous) montane-subalpine |
| *Luzula luzuloides* | kollin-montan (-subalpin) |
| *Luzula sudetica* | montan-subalpin |
| *Lysimachia nummularia* | Kollin-montan (-subalpin) |
| *Malva neglecta* | kollin-montan (-subalpin) |
| *Medicago falcata* | kollin-subalpin |
| *Medicago lupulina* | kollin-montan (-subalpin) |
| *Medicago minima* | kollin-montan |
| Medicago sativa | kollin-montan (-subalpin) |
| Melampyrum pratense | kollin-subalpin (-alpin) |
| Molinia caerulea | kollin-subalpin (-alpin) |
| Myosotis alpestris | (montan-) subalpine-alpine |
| Nardus stricta | montane-alpine |
| Odontites luteus | kollin-montan (-subalpin) |
| Onobrychis sp | kollin-montan (-subalpin) |
| Ononis spinosa | kollin-montan |
| Ophrys sphegodes | kollin (-montan) |
| Orchis mascula | kollin-subalpin (-alpin) |
| Orchis militaris | kollin-montan |
| Orchis ustulata | kollin-subalpin |
| Origanum vulgare | kollin-subalpin |
| Parnassia palustris | kollin-alpin |
| Pastinaca sativa | kollin-montan (-subalpin) |
| Peucedanum oreoselinum | kollin-montan (-subalpin) |
| Phleum pratense | collin-montan (-subalpin) |
| Phyteuma hemisphaericum | (subalpine) alpine |
| Phyteuma orbiculare | montane-subalpine (-alpine) |
| Picea abies | (collinous) montane-subalpine |
| Pimpinella saxifraga | kollin-subalpin |
| Pilosella lactucella | kollin-subalpin |
| Pilosella officinarum | kollin-alpin |
| Plantago alpina | (montane) subalpine-alpine |
| Plantago atrata | (montane) subalpine-alpine |
| Plantago lanceolata | collin-subalpin (-alpin) |
| Plantago major | kollin-subalpin (-alpin) |
| Plantago media | kollin-montan (-subalpin) |
| Platanthera chlorantha | kollin-subalpin |
| Poa alpina | (collinous) subalpine-alpine |
| Poa annua | kollin-subalpin (-alpin) |
| Poa badensis | kollin-montan |
| Poa pratensis | collin-subalpin (-alpin) |
| Polygala alpestris | (montane) subalpine (-alpine) |
| Polygala chamaebuxus | (collinous) montane-subalpine (-alpin) |
| Polygala comosa | kollin-subalpin |
| Polygala vulgaris | kollin-montan (-subalpin) |
| Polygonatum odoratum | kollin-subalpin |
| Polygonum aviculare | kollin-montan (-subalpin) |
| Polygonum viviparum | (montane) subalpine-alpine |
| Potentilla anserina | kollin-montan (-subalpin) |
| Potentilla aurea | (montane) subalpine-alpine |
| Potentilla crantzii | (montane) subalpine-alpine |
| Potentilla erecta | kollin-subalpin (-alpin) |
| Primula auricula | (collin) subalpine-alpine |
| Primula elatior | kollin-subalpin (-alpin) |
| Primula farinosa | (kollin-) montan-alpin |
| Primula integrifolia | (subalpine) alpine |
| Primula veris | Kollin-Subalpin |
| Prunella grandiflora | kollin-subalpin (-alpin) |
| Prunella vulgaris | kollin-subalpin (-alpin) |
| Pteridium aquilinum | kollin-subalpin |
| Pulsatilla alpina subsp. alpina | subalpine-alpine |
| Pulsatilla vernalis | (montane) subalpine-alpine |
| Anemone pulsatilla | kollin-montan |
| Ranunculus repens | kollin-subalpin |
| Ranunculus acris | kollin-subalpin (-alpin) |
| Ranunculus alpestris | (montane) subalpine-alpine |
| Ranunculus bulbosus | kollin-subalpin |
| Ranunculus montanus | (montane) subalpine-alpine |
| Ranunculus tuberosus | kollin-subalpin |
| Reseda lutea | kollin-montan (-subalpin) |
| Rhinanthus alectorolophus | kollin-subalpin (-alpin) |
| Rubus fruticosus | kollin-subalpin |
| Rubus idaeus | collin-subalpin (-alpin) |
| Rubus saxatilis | collin-subalpin (-alpin) |
| Rumex acetosella | kollin-subalpin |
| Rumex alpestris | montane-subalpine (-alpine) |
| Rumex alpinus | (montane) subalpine (-alpine) |
| Salvia pratensis | kollin-montan (-subalpin) |
| Sanguisorba minor | kollin-subalpin |
| Saponaria ocymoides | kollin-subalpin |
| Scabiosa columbaria | Kollin-Montan |
| Scabiosa lucida | (montane) subalpine-alpine |
| Sedum album | kollin-subalpin (-alpin) |
| Selaginella selaginoides | (montane) subalpine-alpine |
| Senecio alpinus | subalpin-alpin |
| Seseli annuum | kollin-montan |
| Sesleria caerulea | (kollin-) montan-alpin |
| Setaria viridis | kollin-montan |
| Silene nutans | kollin-subalpin (-alpin) |
| Silene vulgaris | kollin-subalpin (-alpin) |
| Solanum dulcamara | kollin-montan (-subalpin) |
| Soldanella alpina | (montane) subalpine-alpine |
| Stachys officinalis | kollin-montan |
| Stachys recta | kollin-montan (-subalpin) |
| Stellaria media | kollin-subalpin |
| Taraxacum officinale | kollin-subalpin (-alpin) |
| Taraxacum campylodes | kollin-subalpin (-alpin) |
| Teucrium chamaedrys | kollin-subalpin |
| Teucrium montanum | kollin-subalpin (-alpin) |
| Thalictrum minus | kollin-subalpin (-alpin) |
| Thesium alpinum | montan-alpin |
| Thesium bavarum | kollin-montan |
| Thymus praecox polytrichus | (colline) subalpine-alpine |
| Thymus pulegioides | kollin-subalpin (-alpin) |
| Thymus serpyllum | kollin-subalpin (-alpin) |
| Tofieldia calyculata | (kollin-) montan-alpin |
| Tragopogon pratensis | kollin-montan (-subalpin) |
| Trifolium aureum | kollin-montan (-subalpin) |
| Trifolium badium | subalpine (-alpine) |
| Trifolium montanum | kollin-subalpin (-alpin) |
| Trifolium pratense | kollin-subalpin (-alpin) |
| Trifolium repens | kollin-subalpin (-alpin) |
| Trifolium thalii | subalpine-alpine |
| Trisetum flavescens | Kollin-montan (-subalpin) |
| Trollius europaeus | (collin-) montane-subalpin (-alpin) |
| Tussilago farfara | kollin-subalpin (-alpin) |
| Urtica dioica | kollin-subalpin (-alpin) |
| Vaccinium myrtillus | kollin-alpin |
| Vaccinium vitis-idaea | (Kollin-) montan-subalpin |
| Valeriana montana | (montane) subalpine (-alpine) |
| Valerianella locusta | kollin-montan |
| Veratrum album | montan-alpin |
| Verbena officinalis | kollin-montan (-subalpin) |
| Veronica aphylla | (montan-) subalpine-alpine |
| Veronica chamaedrys | kollin-subalpin (-alpin) |
| Veronica fruticans | (montan-) subalpine-alpine |
| Veronica officinalis | kollin-subalpin |
| Veronica persica | kollin-montan (-subalpin) |
| Veronica spicata | kollin-subalpin |
| Veronica teucrium | kollin-montan (-subalpin) |
| Vicia cracca | kollin-subalpin |
| Vicia pratensis | kollin-montan (-subalpin) |
| Vicia sepium | kollin-montan (-subalpin) |
| Vincetoxicum hirundinaria | kollin-subalpin |
| Viola calcarata | subalpine-alpine |
| Viola hirta | kollin-montan (-subalpin) |
| Viola tricolor | (collinous) montane-subalpine |

**Table S3.** Comparison of different models quantifying the relationship between host community traits and disease. Each model contained square-root transformed community parasite load as the response, and elevation, host community species richness, richness-independent phylogenetic diversity, and some combination of host traits as fixed effects. To estimate whether the effect of host community structure depends on elevation, we also included in the model the pairwise interactions between each measure of host community structure and elevation as additional fixed effects, The Pace-of-Life model includes host community pace of life as a latent factor, and is the model reported in the manuscript. The All Traits model includes all single traits in place of the pace of life latent factor. The Chlorophyll, Leaf Longevity, Leaf Nitrogen, Leaf Phosphorus, and Specific Leaf Area models include a single trait in place of the latent factor.

| Model | AICc | Marginal<br>R<sup>2</sup> | Conditional<br>R<sup>2</sup> | RMSE | LOOCV<br>RMSE |
| --- | --- | --- | --- | --- | --- |
| Pace-of-Life | 207.99 | 0.238 | 0.46 | 0.295 | 0.315 |
| All Traits | 319.42 | 0.252 | 0.432 | 0.293 | 0.319 |
| Chlorophyll | 220.28 | 0.167 | 0.393 | 0.296 | 0.318 |
| Leaf Longevity | 217.93 | 0.183 | 0.434 | 0.292 | 0.315 |
| Leaf Nitrogen | 221.72 | 0.166 | 0.393 | 0.294 | 0.317 |
| Leaf Phosphorus | 220.09 | 0.176 | 0.399 | 0.298 | 0.318 |
| Specific Leaf<br>Area | 212.41 | 0.251 | 0.391 | 0.298 | 0.316 |

**Table S4.**
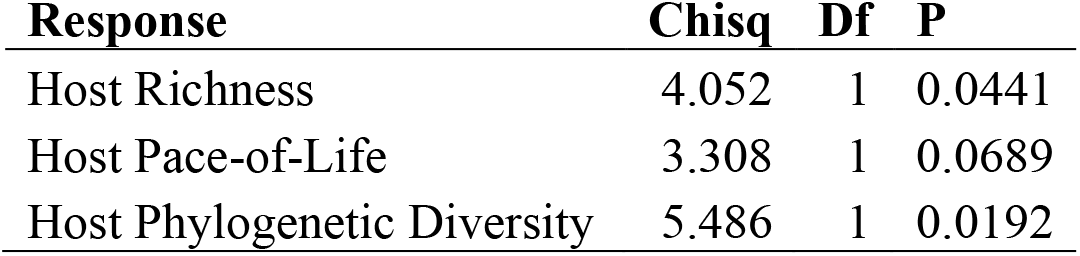
Results of Type II Analysis of Deviance tests on models quantifying the effects of elevation on three measures of host community structure.

| Response | Chisq | Df | P |
| --- | --- | --- | --- |
| Host Richness | 4.052 | 1 | 0.0441 |
| Host Pace-of-Life | 3.308 | 1 | 0.0689 |
| Host Phylogenetic Diversity | 5.486 | 1 | 0.0192 |

**Table S5.** Results of Type II Analysis of Deviance test on the mixed model of disease, testing whether each factor influenced square-root transformed community parasite load.

| Predictor | Chisq | Df | P |
| --- | --- | --- | --- |
| Elevation | 7.0917 | 1 | 0.0077 |
| Host Richness | 7.6565 | 1 | 0.0057 |
| Host Phylogenetic Diversity | 0.1950 | 1 | 0.6588 |
| Host Pace-of-Life | 1.7552 | 1 | 0.1852 |
| Elevation × Host Richness | 1.5493 | 1 | 0.2132 |
| Elevation × Host Phylogenetic Diversity | 1.0018 | 1 | 0.3169 |
| Elevation × Host Pace-of-Life | 9.9692 | 1 | 0.0016 |

**Table S6.** Coefficient estimates from the structural equation model. Estimates are provided both raw and standardized to a common scale to facilitate comparisons. Correlations among dependent variables are indicated by ~~.

| Response | Predictor | Estimate | Std Error | DF | Critical Value | P | Std Estimate |
| --- | --- | --- | --- | --- | --- | --- | --- |
| Square-root transformed community parasite load | Elevation | -0.0002 | 0.0001 | 86 | -3.598 | 0.0005 | -0.2315 |
|  | Host Species Richness | -0.0107 | 0.0039 | 106 | -2.727 | 0.0075 | -0.1557 |
|  | Host Phylogenetic Diversity | -0.0106 | 0.0231 | 106 | -0.460 | 0.6466 | 0.0244 |
|  | Host Pace-of-Life | 0.7397 | 0.2104 | 106 | 3.515 | 0.0006 | 0.6551 |
| | Elevation × Pace-of-Life | -0.0005 | 0.0002 | 106 | -3.258 | 0.0015 | -0.6104 |
| Host Species Richness | Elevation | 0.0031 | 0.0016 | 86 | 2.013 | 0.0472 | 0.2443 |
| Host Pace-of-Life | Elevation | -0.0002 | 0.0001 | 86 | -1.819 | 0.0724 | -0.3019 |
| Host Phylogenetic Diversity | Elevation | 0.0003 | 0.0001 | 86 | 2.342 | 0.0215 | 0.1595 |
| ∼ Host Species Richness | ∼ Host Pace-of-Life | -0.0871 | NA | 220 | -1.288 | 0.0996 | -0.0871 |
| ∼ Host Phylogenetic Diversity | ∼ Host Pace-of-Life | -0.0775 | NA | 220 | -1.145 | 0.1268 | -0.0775 |

Goodness of fit: Fisher's C = 1.463 with p = 0.481 and on 2 degrees of freedom

## Figure Legends

**Figure S1.**
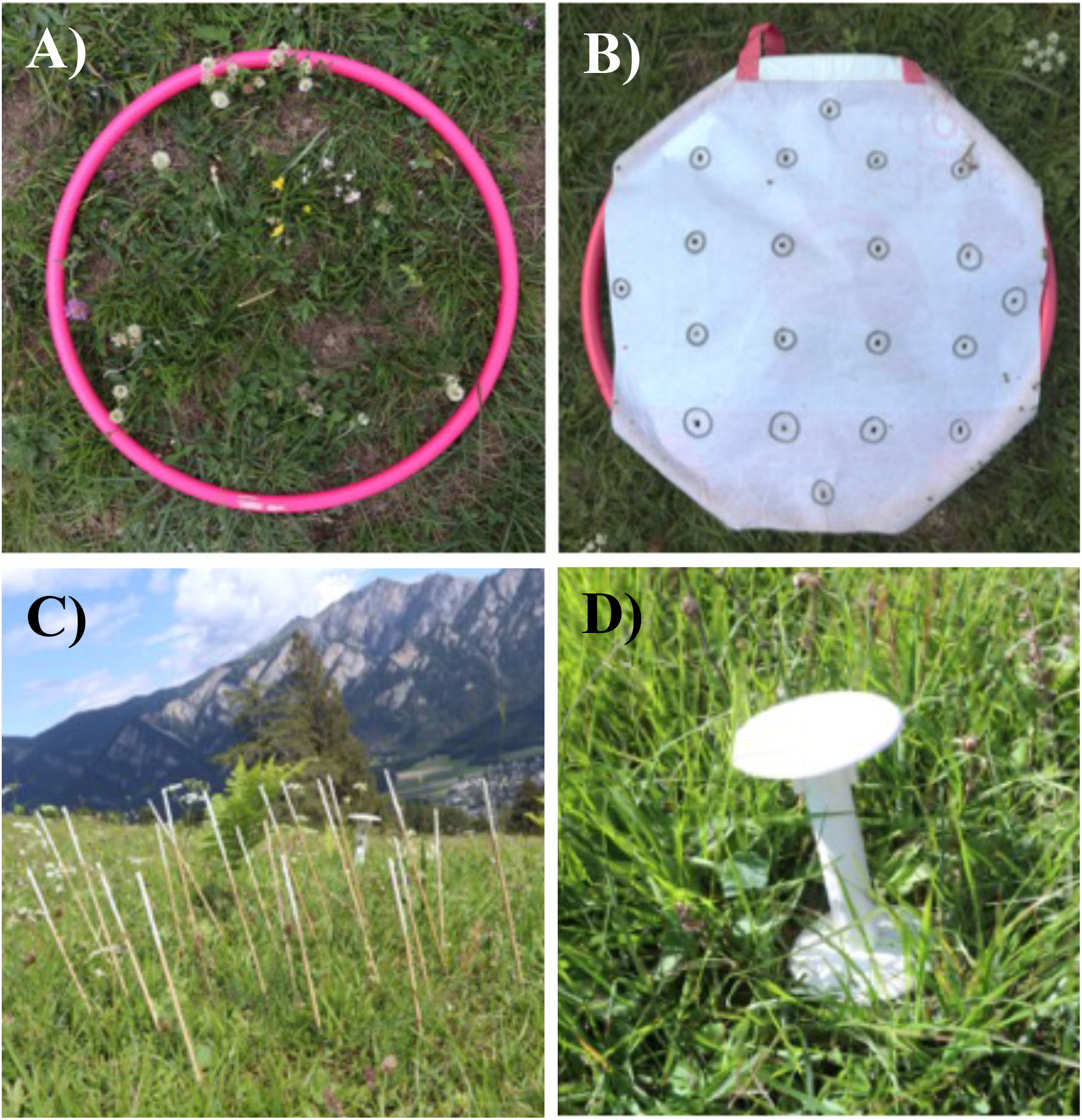
Images representing survey methods. A) The vegetation survey was carried out in small plots that were marked with a central stick and delineated by a 50 cm-diameter hula hoop. The disease survey was carried out by B) placing 20 grill sicks through a 50 cm-diameter hula hoop covered with canvas containing 20 evenly spaced holes and C) surveying the plant most touching each grill stick for foliar disease symptoms. D) Air temperature, soil surface temperature, soil temperature, and soil moisture were recorded in 15-minute intervals using TOMST-4 data loggers that were placed in the central large plot of each site. Photos: Mikko Jalo

**Figure S2.**
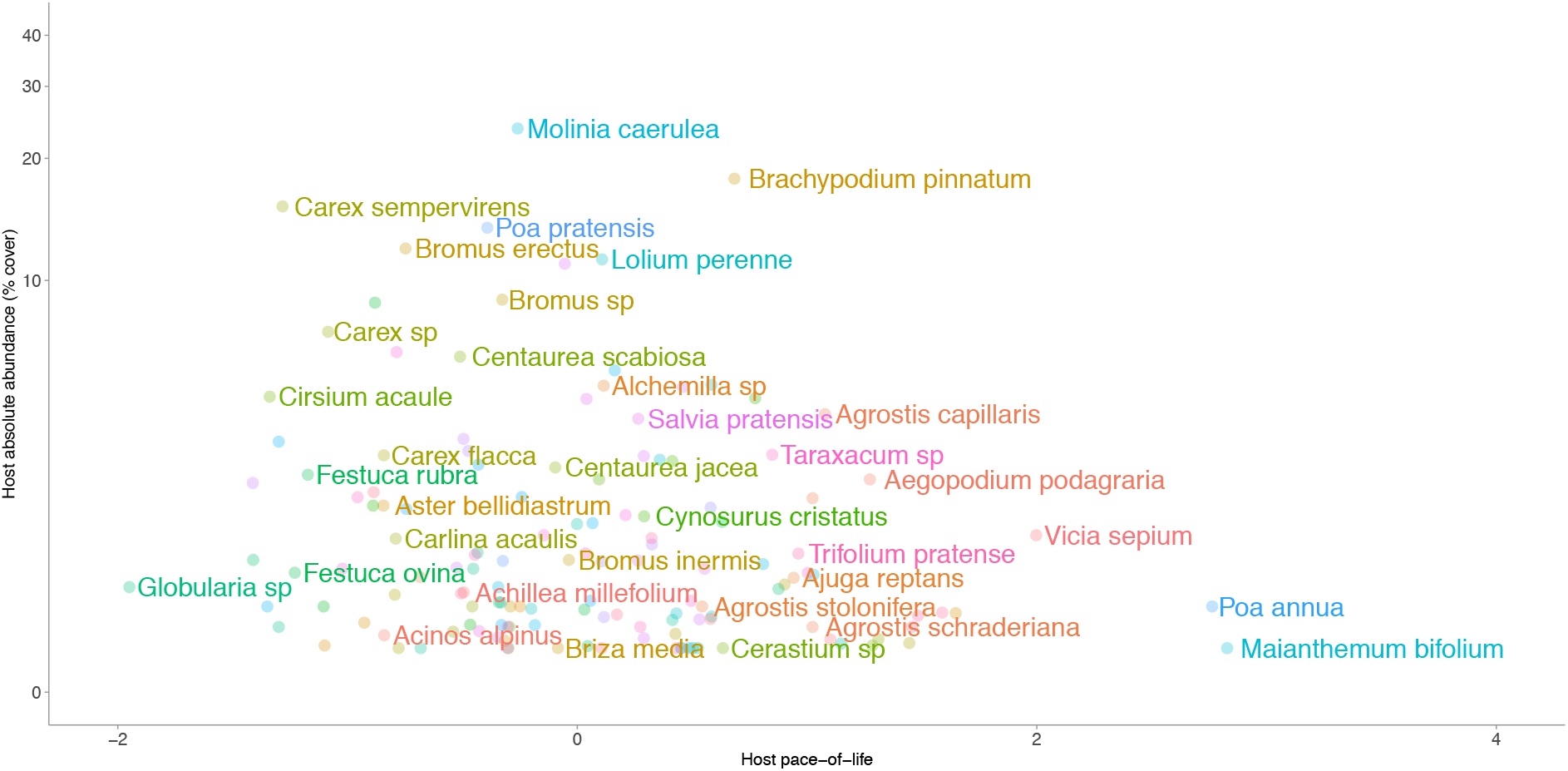
Host taxa arranged by their mean absolute vegetative cover in plots where those taxa occur (y axis) and host pace of life (x-axis). Increasing values of host-pace-of life are associated with increases in specific leaf area, leaf chlorophyll content, leaf nitrogen, and leaf phosphorus, and with shorter leaf lifespans. Taxa with the highest local abundance tended to exhibit intermediate life-history strategies, while taxa exhibiting extreme life-history strategies tended to be locally rare.

**Figure S3.**
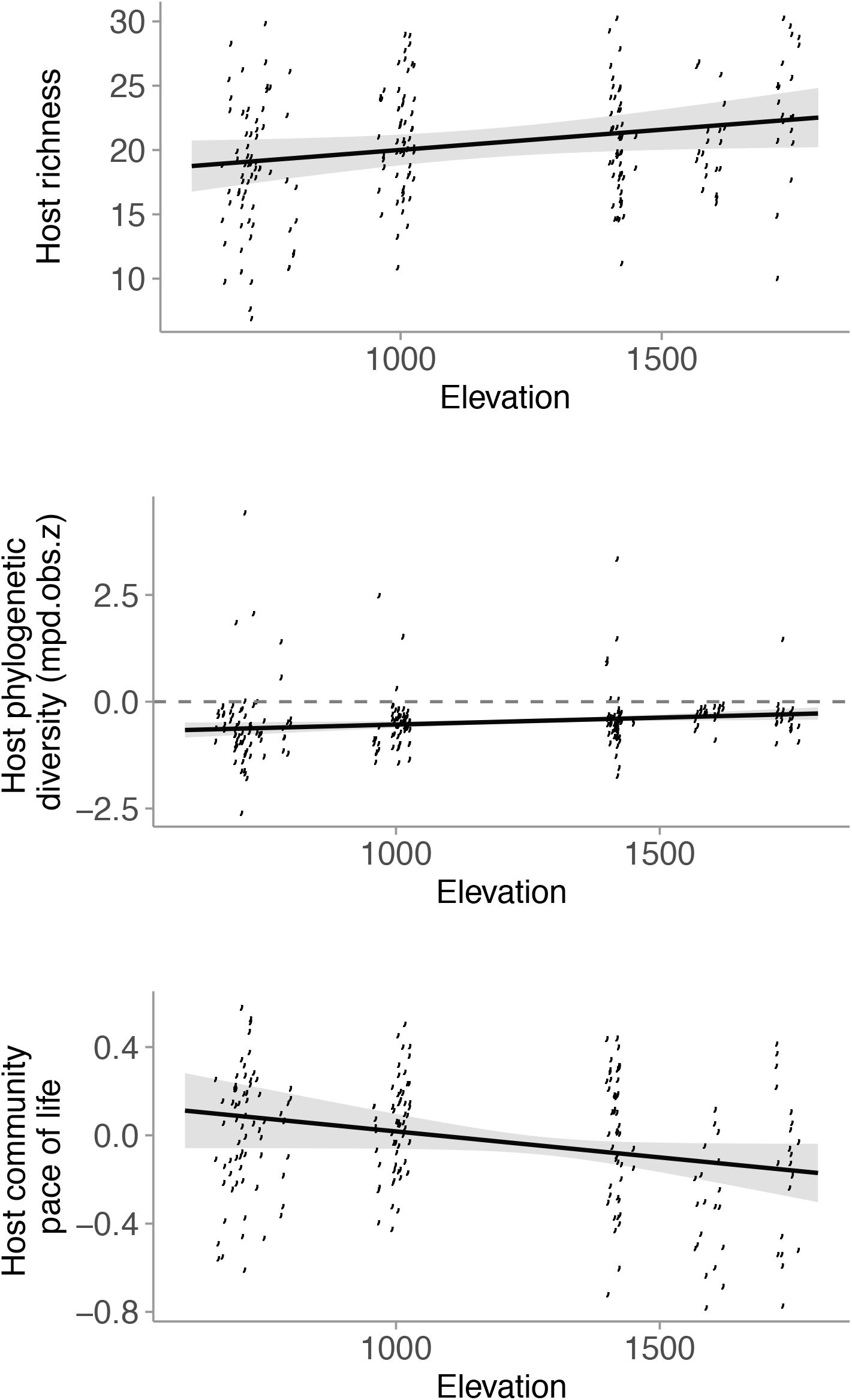
Relationship between host richness, richness-independent host phylogenetic diversity, and host community pace of life (together measuring host community structure) and elevation. Species richness increased with increasing elevation. Host communities tended to be more phylogenetically clustered than random, and the degree of phylogenetic clustering decreased (i.e., phylogenetic diversity increased) with increasing elevation. Host community pace of life declined slightly with increasing elevation, though this relationship was not supported by statistical models.

